# deepMINE - Natural Language Processing based Automatic Literature Mining and Research Summarization for Early-Stage Comprehension in Pandemic Situations specifically for COVID-19

**DOI:** 10.1101/2020.03.30.014555

**Authors:** Bhrugesh Joshi, Vishvajit Bakarola, Parth Shah, Ramar Krishnamurthy

## Abstract

The recent pandemic created due to Novel Coronavirus (nCOV-2019) from Wuhan, China demanding a large scale of a general health emergency. This demands novel research on the vaccine to fight against this pandemic situation, re-purposing of the existing drugs, phylogenetic analysis to identify the origin and determine the similarity with other known viruses, etc. The very preliminary task from the research community is to analyze the wide verities of existing related research articles, which is very much time-consuming in such situations where each minute counts for saving hundreds of human lives. The entire manual processing is even lower down the efficiency in mining the information. We have developed a complete automatic literature mining system that delivers efficient and fast mining from existing biomedical literature databases. With the help of modern-day deep learning algorithms, our system also delivers a summarization of important research articles that provides ease and fast comprehension of critical research articles. The system is currently scanning nearly 1,46,115,136 English words from 29,315 research articles in not greater than 1.5 seconds with multiple search keywords. Our research article presents the criticality of literature mining, especially in pandemic situations with the implementation and online deployment of the system.

## 1 Introduction

The recent pandemic of coronavirus disease (COVID-19)[1] is caused by novel human coronavirus, initially referred to as the Wuhan coronavirus (CoV), which is currently designated as a severe acute respiratory syndrome (SARS)-CoV-2 as per the latest International Committee on Taxonomy of Viruses (ICTV) classification[2] and was suggested to have a possible zoonotic origin[3]. As an outbreak of pneumonia started in Wuhan, China and the first case was found on 12 December 2020. By the end of March 2020, it has infected more than 750,000 individuals in nearly 201 countries and causes more than 36,000 deaths worldwide[4].

SARS-CoV-2 belongs to the family of Coronaviridae. It has an envelope and maintains a single-strand, positive-sense RNA genome ranging from 26 to 32 kb in length[5]. These viruses can be classified into four genera: alpha, beta, delta, and gamma. From which alpha and beta Coronavirus (CoVs) are known to infect humans[6]. They are circulated among humans, other mammals, and birds. They can cause respiratory, hepatic, enteric, and neurologic diseases[7] as well. Despite the fact majority of human coronavirus infections found to have a mild effect, the epidemics of two Beta coronaviruses (betaCoV) - namely Severe Acute Respiratory Syndrome Coronavirus (SARS-CoV) and the Middle East Respiratory Syndrome Coronavirus (MERS-CoV) have caused more than ten thousand cumulative cases in the past 20 years with mortality rates about 10 percent for SARS-CoV and 37 percent in case of MERS-CoV [8][9].

As the situation is getting worst turning from epidemic to pandemic with an increasing number of confirmed cases and its related deaths with each passing day. Science community around the world has joined their hand to fight against this deadly disease in all the possible ways including, making the vaccines, repurposing the existing drugs and designing diagnostic kits for the detection of the presence of virus or disease. Varieties of works also includes sequencing the virus genome to identify the origin of the virus and the possible mode of transmission, to allocate the resource for the patients and applying the statistical models to predict how fast the disease can be spread.

To begin any new research it is very essential to know what information is already available. As we know the fact that the similar virus has already been known, it becomes crucial to collect and organize this information from the ocean of research articles. The average reading speed of a human is roughly 200 wpm (words per minute)[10], that means that it will take a substantial amount of time to comprehend and find useful information from thousands of research articles published. To help the research community for screening the thousands of published research and making the comprehension in a short amount of time we have designed the AI-NLP based system which can mine the relevant articles and give a short article summary, that can make ease in fast and efficient comprehension for researchers. As various experts are working in a different domain they search for the relevant information which lies in a varied domain.

Data mining is becoming a fundamental component of the global world with verities of applications. Data mining assists a quick and efficient decision-making process that enhancing the accuracy of an outcome. We already have a huge amount of data or information in forms of research articles published in the last 5 decades or more. With the application of the literature mining system, meaningful research articles can be extracted from this huge set and brief technical summary can be produced for research articles user interested with.

In computer science, text summarization is a process of shortening the large text document(s) in order to generate short and meaningful piece of text. The objective is to create fluent natural language text keeping major insights or technicality of the source data. The automatic text summarization is an ordinary problem in the field of natural language processing and machine learning. The task was first carried out in form of generating automatic literature abstracts in 1958 [11]. Over the past half a century, the problem of text summarization has been addressed with verities of perspectives.

Primarily, the task of summarization is divided into two major categories as *Extractive summarization* and *Abstractive summarization* [12]. As the name itself suggests, the Extractive method involves pulling of key phrases from the source document in order to generate the targeted summary. The Abstractive summarization works similar as we human do [13]. It involves the end-to-end deep learning technique called Sequence-to-Sequence learning to derive the understanding about the association between words.

The primary objective of the our system is to deliver quick and efficient search from a huge amount of available literature. By entering a keyword *CORONAVIRUS* we are getting 2116 number of research articles that contain the keyword in the title. This is even time consuming to go through the abstract of each literature, here we are interested in. Hence we found that searching for an article is not serves the ultimate purpose of serving important research articles to a user so that the researcher can speed up the research in epidemic situations. To overcome this limitation we developed a research text summarizer that can generate a technical summary by scanning all the research articles derived from user-entered keyword(s). In the demanding situation of COVID-19, we applied the literature mining with user entered keyword(s) and automatic generation of brief summary of research articles, that user searches for. The ultimate objective of our system deepMINE, is to provide quick and efficient access of the openly available research articles.

## 2 Dataset Used

For the initial starting purpose, we have used the COVID-19 Open Research Dataset (CORD-19) [14]. The dataset has been published by Allen Institute for AI on 20th March 2020. The detailed category wise description of the CORD-19 dataset is given in Table-1.

**Table 1.** Category wise description of CORD-19 dataset.

| Sr. No. | CORD-19 Category | Number of Articles | Number of Words |
| --- | --- | --- | --- |
| 1 | Commercial use subset | 9118 | 43,605,042 |
| 2 | Non-commercial use subset | 2352 | 10,165,054 |
| 3 | Custom license subset | 16959 | 89,279,050 |
| 4 | bioRxiv/medRxiv pre-prints subset | 885 | 3,065,990 |
| **Total** |  | **29315** | **1,46,115,136** |

## 3 Implementation

The deepMINE is primarily performing two major functions namely mining of articles from available open data sources using user-entered keywords and generate brief technical summary in natural language for a quick review of articles that user interested with. The entire system has been developed using Python programming language with the support of various scientific and natural language processing libraries available.

As represented in Figure-1, a researcher can submit the keyword(s) he or she is interested with and the **Mine Article** process of the system returns article titles and available links by scanning more than 1,46,115,136 words from 29,315 research articles openly provided by CORD-19. A user of the system can separately make requests for generating a summary of the research articles with **Article Summarization** process. The system has used the deep natural language processing-based text summarization for generating detailed technical summary given the research article as an input.

**Fig. 1.**
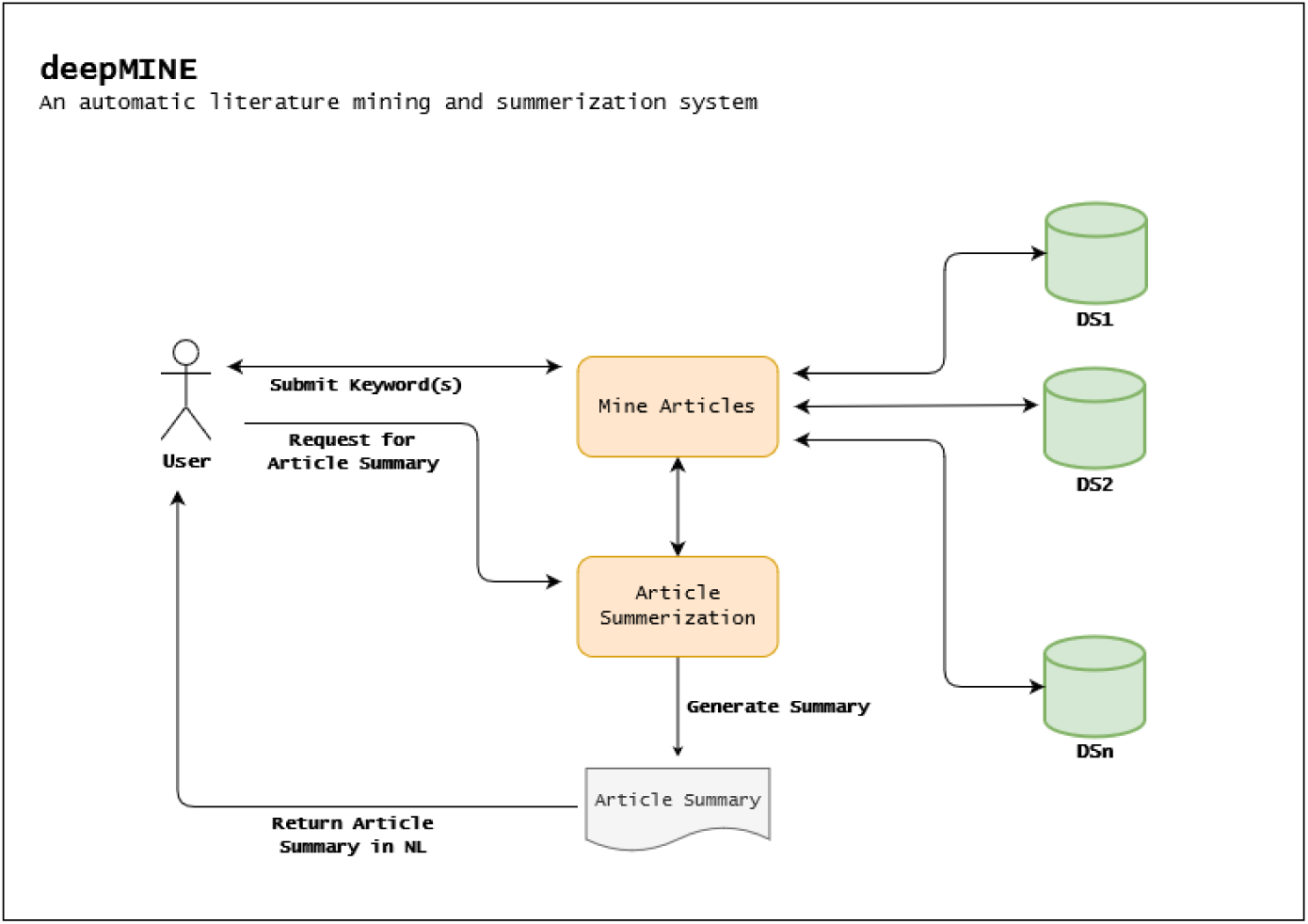
Data flow diagram of deepMINE.

## 4 Experiments and Results

The deepMINE system uses the natural language processing based text mining from available data sources. One user enters keyword(s) related to articles he or she is interested with, system returns all the articles having the entered keyword(s) in the article title. For returning more precise articles, we have currently focused on the title of the available research articles.

The system returns found research articles having the keyword(s) in the title with total count and the user can explicitly visit to abstract and summary to the research article. The deepMINE service is openly available on URL https://deepmine.in/

As discussed under Section - 2, CORD-19 dataset contains nearly 1,46,115,136 words in *title, abstract* and *article body*. The below Table-2 represents the dataset category wise title word count.

**Table 2.** Category wise title words count in CORD-19.

| Sr. No. | CORD-19 Category | Number of Articles | Number of Words in Title |
| --- | --- | --- | --- |
| 1 | Commercial use subset | 9118 | 129,148 |
| 2 | Non-commercial use subset | 2352 | 31,188 |
| 3 | Custom license subset | 16959 | 190,250 |
| 4 | bioRxiv/medRxiv pre-prints subset | 885 | 12,695 |
| **Total** |  | **29315** | **363,281** |

While mining through the deepMINE given a keyword “CORONAVIRUS” we found 2116 number of research articles as shown in Figure-2.

**Fig. 2.**
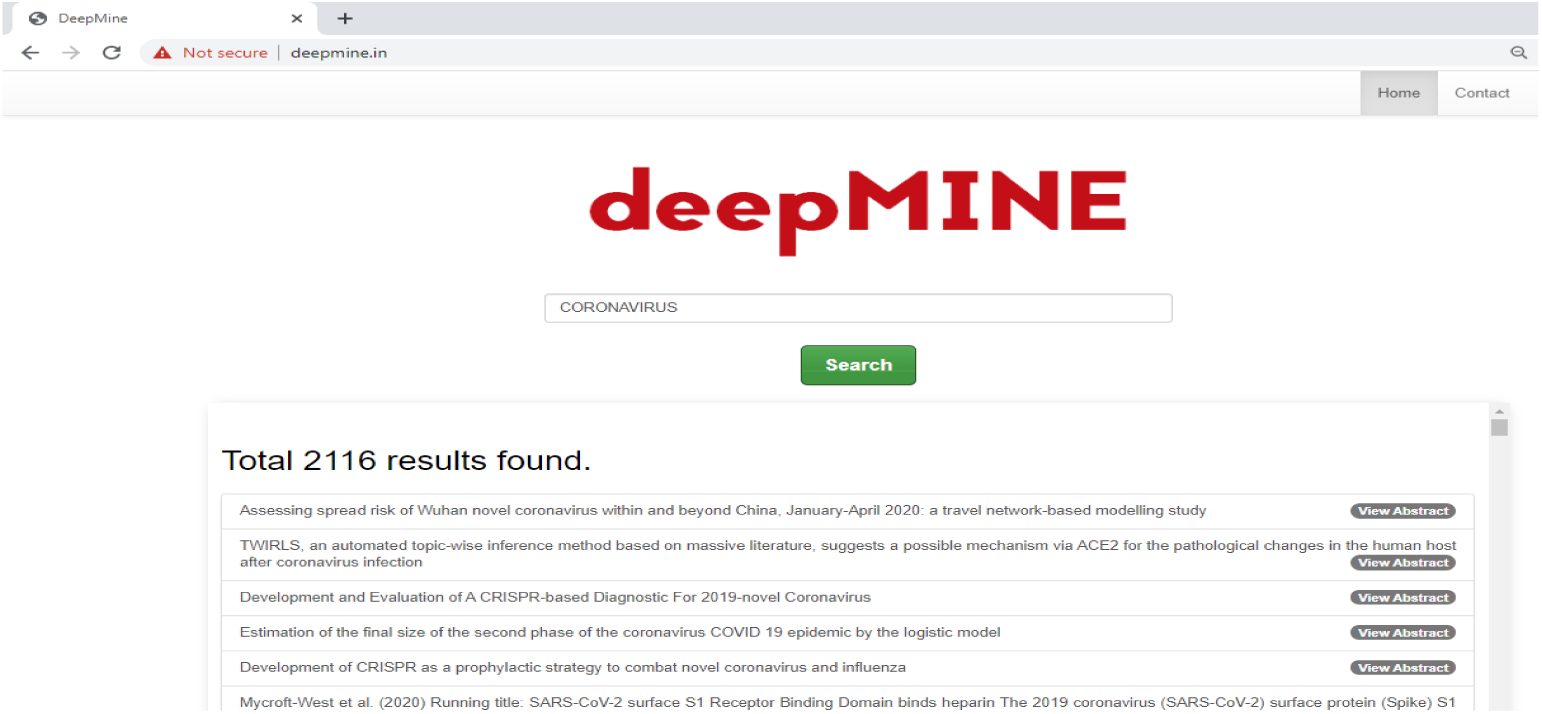
Result of literature mining with keyword “CORONAVIRUS”.

Among total 2116 number of research articles mined using keyword “CORONAVIRUS”, Figure-3 demonstrate country wise article research article count with given keyword.

**Fig. 3.**
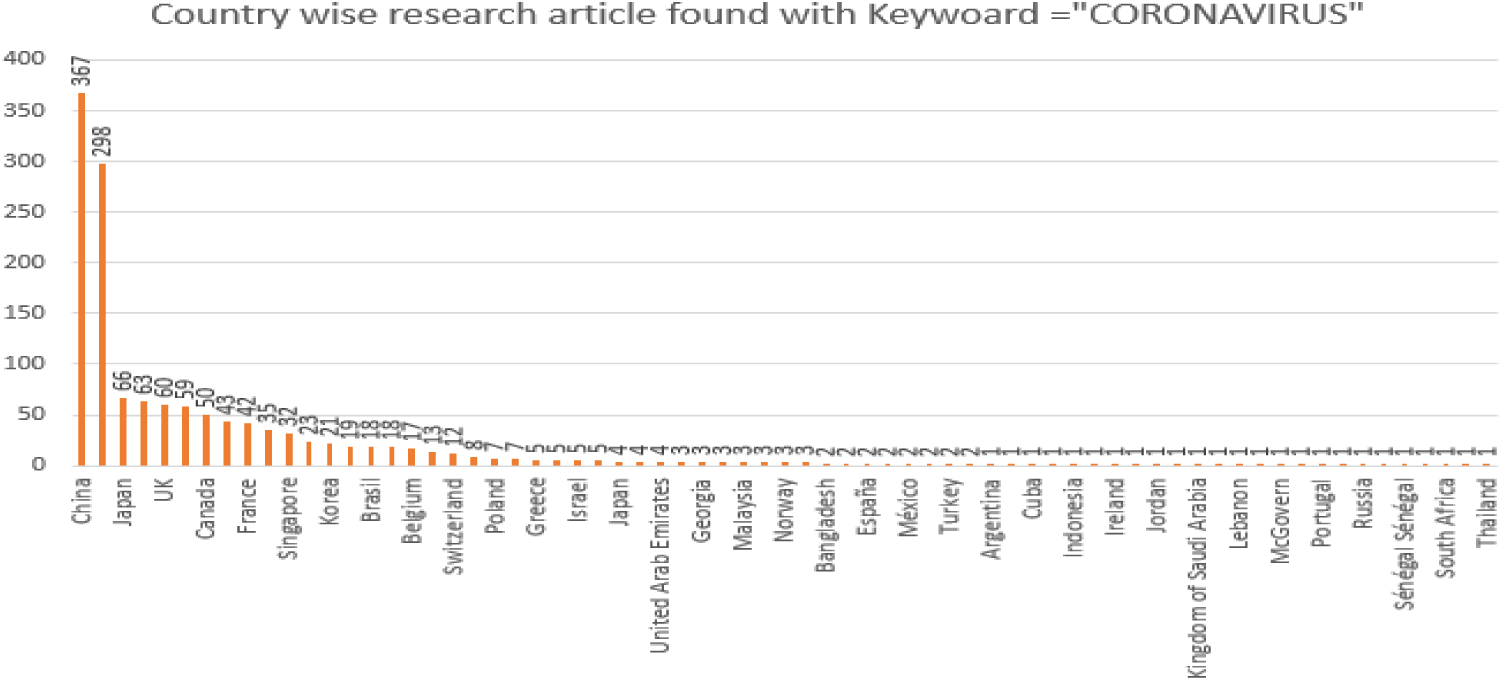
Graph showing country wise research articles mined using keyword “CORONAVIRUS”.

It is utmost significance for a researcher to take a quick idea of research with the abstract of the research article. deepMINE system derived abstract for all mined articles from the dataset for a quick review. The below Figure-4 shows the abstarct of one of the research articles mined suing keyword “CORONAVIRUS”.

**Fig. 4.**
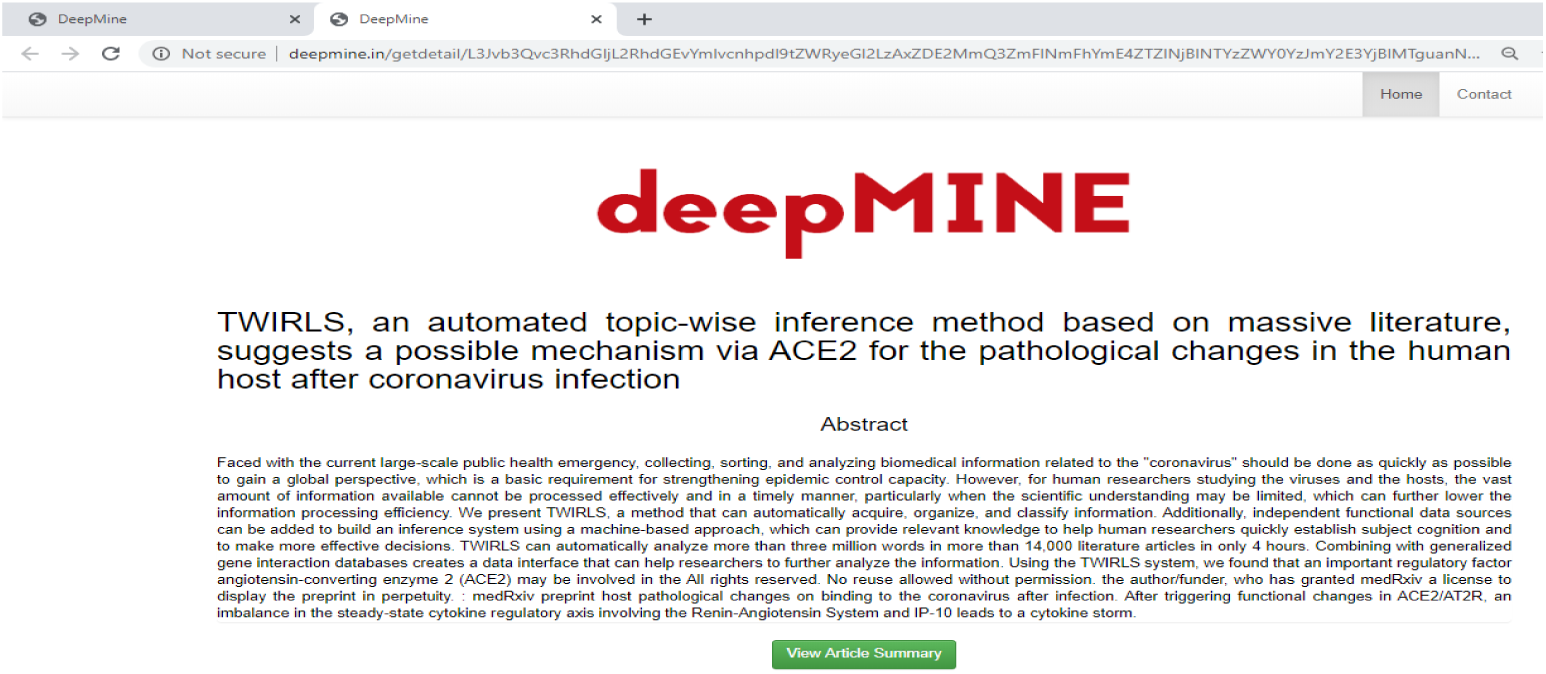
Absract view of one of the articles found with keyword “CORONAVIRUS”.

From the homepage of the system as shown in Figure-2, we mined research articles using multiple keywords i.e. “CORONAVIRUS” and “RNA”. This mining returns us total 98 results. Statistics of country wise research articles as shown in Figure-5 and abstract of one of the research articles is shown in Figure-6.

**Fig. 5.**
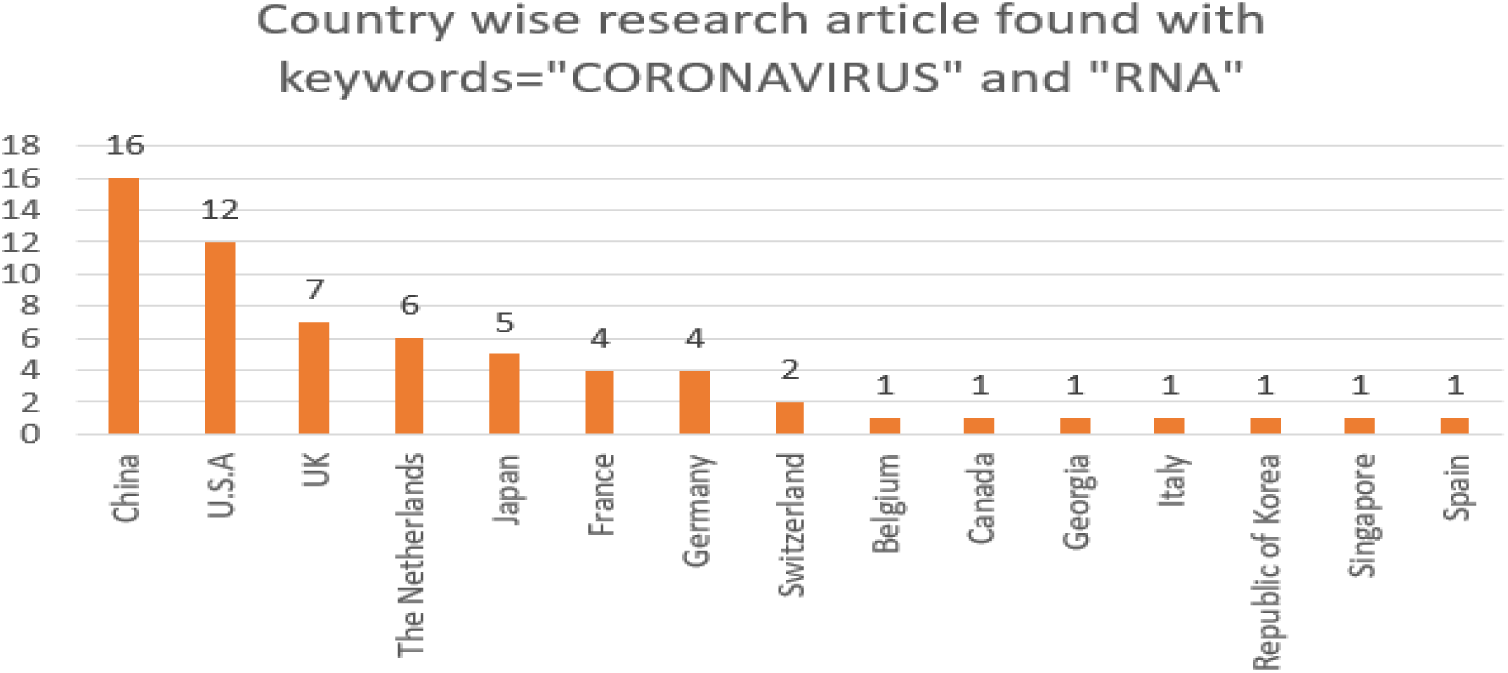
Graph showing country wise research articles mined using keywords “CORONAVIRUS” and “RNA”.

**Fig. 6.**
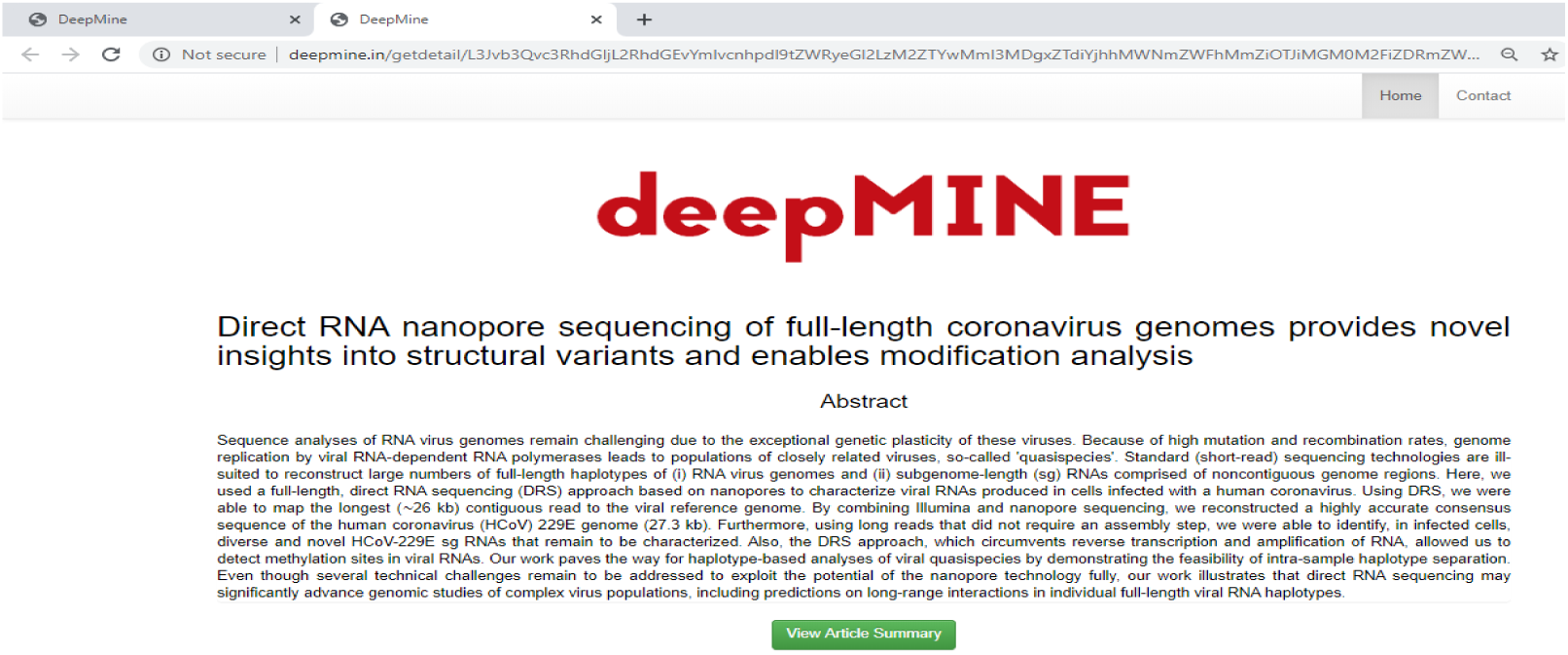
Absract view of one of the articles found with keywords “CORONAVIRUS” and “RNA”.

## 5 Conclusion

Our primary contribution majorly focuses on the design and development of the system that can able to automatically mine the literature, provided the keyword(s) as an input. The system is able to generate the brief summary of the research article, that user interested with. Our ultimate objective is to provide a quick and easy literature mining service so that the researchers working for fighting current pandemic situation of COVID-19 can get early-stage comprehension. Currently we have used the COVID-19 Open Research Dataset (CORD-19), publicly made available by Allen Institute of AI on 20th March 2020. Our system deepMINE is providing mining from 29,315 research articles with keywords by scanning nearly 1,46,115,136 English words available in literature dataset in not greater than 1.5 seconds.

From experiments performed on the live deepMINE system we can observer that the system is helpful for the primary mining of the huge amount literature available in various openly available datasets. However, in many of the research articles of the dataset the affiliation and/or location has not been provided. We are in the process of cleaning and improving the dataset with automatic and manual processes.

## 6 Future work plan

Currently, we are working on the collection of various research articles openly available from reputed publishers. Afterward, the system will be upgraded with a wide range of research literature especially of the bio-medical field.

An equal amount of work is going on to improve the process of text summarization with the latest deep learning techniques to provide more accurate and human like text summarization on research articles.

